# A portable, low-cost device for precise control of specimen temperature under stereomicroscopes

**DOI:** 10.1101/2019.12.19.882605

**Authors:** Nicholas D. Testa, Samiksha Kaul, Kim N. Le, Mei Zhan, Hang Lu, Annalise B. Paaby

## Abstract

To facilitate precise and convenient control of biological sample temperature, we developed a low-cost device that can be used independently or with any stereomicroscope. The purpose of the device is to control the thermal environment during experimental intervals in which a specimen must be manipulated outside of an incubator, e.g. for dissection or slide-mounting in preparation for imaging. Sample temperatures can be both cooled to below and heated to above room temperatures, and stably maintained at a precision of +/− 0.1°C. To demonstrate the utility of this device, we report improved characterization of the penetrance of a short-acting temperature-sensitive allele in *C. elegans* embryos, and identification of the upper temperature threshold for embryonic viability for six *Caenorhabditis* species. By controlling the temperature environment even as a specimen is manipulated, this device offers consistency and flexibility, reduces environmental noise, and enables precision timing in experiments requiring temperature shifts.

## INTRODUCTION

The function of living systems depends intimately on temperature. For example, the development of virtually all poikilotherms is sensitive to temperature, even for wild type animals in a normal thermal range [1]. Developmental timing in particular is highly responsive to temperature [2, 3]; the fruit fly *Drosophila melanogaster* and the nematode worm *Caenorhabditis elegans* develop twice as fast at 25°C compared to 18°C and 16°C, respectively [4, 5], and the rate of nematode embryonic cell divisions increases exponentially with temperature [6]. At the mechanistic level, temperature affects molecular events such as gene expression and protein stability [7, 8]. Investigative research often depends on temperature manipulation as part of the experimental process, for example the induction of heat shock or the use of temperature-sensitive mutant alleles. Consequently, experiments involving living samples will be optimized by precise control of the thermal environment. In some cases, engineering near-complete and automated control over the thermal environment has allowed new access into testing hypotheses about environmental variables, for example on health and lifespan [9].

To facilitate precise and convenient control of biological sample temperature, we developed a portable device that can be used independently or with any stereomicroscope. Its purpose is to control the thermal environment during experimental intervals in which a specimen must be manipulated outside of an incubator, e.g. for dissection or slide-mounting in preparation for imaging, or for observational data collection. It is a stand-alone unit, requiring a conventional 120 volt electrical outlet but no computer. Sample temperatures can be both cooled to below and heated to above room temperatures, and stably maintained at a precision of +/− 0.1°C. Our specifications include a flat platform for slides or coverslips and a recessed platform for small petri plates, such as those used in *C. elegans* research. Samples on the device can be illuminated from above or obliquely, ideally through flexible fiber-optic goosenecks. In this configuration, the etched mirror platform reflects light back through the sample, accommodating samples that are typically illuminated from below by a transmitted light base.

By controlling the temperature environment even as a specimen is manipulated, this device offers consistency and flexibility, reduces environmental noise, and enables precision timing in experiments requiring temperature shifts. To demonstrate this utility, we conducted two types of experiments. First, we used temperature to induce an engineered molecular response, and we report improved penetrance and better characterization of a short-acting temperature-sensitive allele in *C. elegans* embryos. Second, we used precise control of temperature to identify naturally-occurring variation in thermal tolerance, and we report identification of the upper temperature threshold for embryonic viability for six *Caenorhabditis* species.

## MATERIAL AND METHODS

### Device components and functionality

#### Core assembly

The core device is comprised of a thermoelectric cooler (TEC), which uses electrical energy to transfer heat from one side to the other; a solid copper heatsink affixed to one side of the TEC, which provides a working surface with even heat distribution to the specimen; a fan-controlled CPU heatsink affixed to the other side of the TEC, which dissipates heat; a thermistor, which reads temperature from the specimen or the working surface of the device; and a microcontroller board and two additional shields (bi-directional motor driver board and protoboard), which control the electrical draw required by the TEC to maintain the desired temperature (Fig 1). Additional components described herein, including buttons and a liquid crystal display (LCD) screen, add functionality and enable real-time feedback of temperature data, but are not required for operation. The components we used to build the device, and their sources and prices, are listed in File S1.

**Fig 1.**
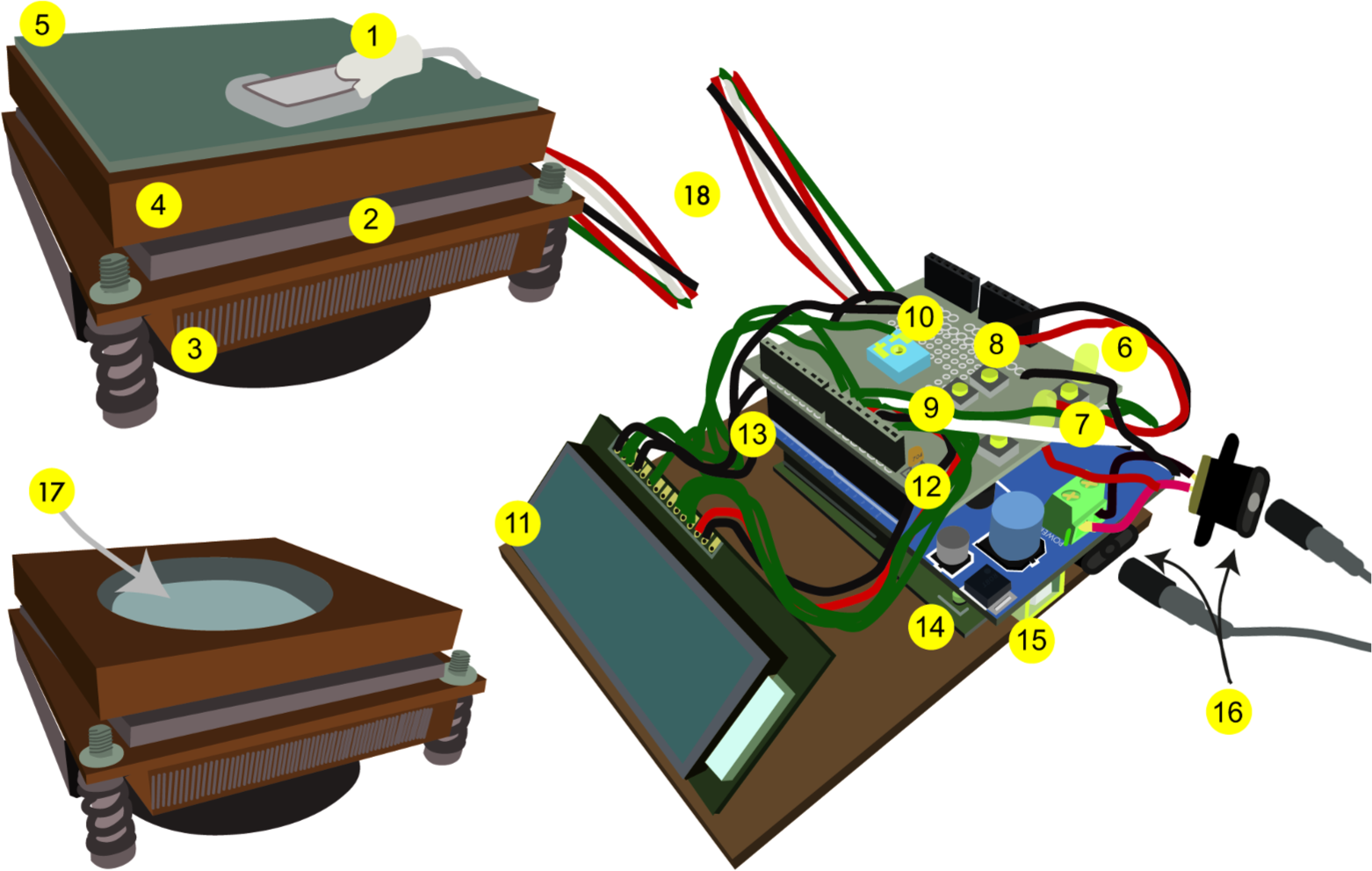
Device overview. (1) Thermistor. (2) Thermoelectric cooler (TEC). (3) Fan-controlled CPU heatsink. (4) Solid copper heatsink. (5) Mirror. (6) PID indicator light, which increases in intensity as more energy is drawn to heat or cool the stage. (7) Reset button. (8) Setpoint temperature + button. (9) Setpoint temperature – button. (10) Potentiometer, which adjusts the intensity of the LCD screen. (11) LCD screen. (12) Protoshield (top board). (13) Bi-directional motor driver shield (middle board). (14) Arduino microcontroller (bottom board). (15) USB port. (16) Power supply ports and cables; the two power supplies can be plugged into a power strip (not shown), and the device powered on or off using the power strip switch. (17) Alternate heatsink with a well for a petri plate. (18) Wires connecting the microcontroller unit and the heatsink unit on which the specimen is mounted can be of any length.

As designed, the heating/cooling component of the device, which includes the working surface for the biological sample, is separate from the microcontroller and connected by wires of any desirable length; this modular design allows the sample to be moved freely, and for easy substitution of working surfaces with different specifications. For example, we used a flat copper slab as a working surface for specimens mounted on slides, and one with a drilled recess for samples on petri plates (Fig 1). As such, the thermistor can be mounted on the working surface of the device, to measure temperature as experienced by specimens on coverslips or slides, or can be embedded in a sample environment, such as agar medium within a plate. While we initially prototyped circuit design with a breadboard, we do not recommend this for long-term use due to the high current demands of the TEC and the somewhat fragile nature of breadboard circuits. Instead, all wires are soldered onto a protoboard shield. The entire assembly is therefore durable, lightweight, versatile, and portable.

#### Arduino microcontroller board

The Arduino Uno board (Arduino.cc, Italy) is an open source platform based on a low power 8-bit microcontroller (Atmega328; Atmel Corp., San Jose, CA) with 32K bytes in programmable flash memory. The board (16 MHz) has 14 digital I/O (Input/Output) pins, and 6 analog input pins. A USB port both powers the board and functions as a communication terminal for data transfer with a computer, when necessary.

#### Bi-directional motor driver board

The Arduino board communicates with sensors via 5.0V I/O pins, while the TEC’s optimal functioning range is 14V/10A. Thus, we employed a bi-directional motor driver shield and external power supply (12V/10A) to amplify the signals communicated by the Arduino board to the TEC. A bi-directional motor driver has many distinct advantages over a traditional MOSFET or Transistor style, specifically in its ability to reverse the flow of electricity to the TEC. Since the flow of electricity within the TEC determines the direction of heat transfer, being able to switch directions allows us to heat or cool the specimen on the same surface.

#### Prototype board and circuit

Due to the fragile nature of breadboard circuits and their low voltage/current limitations, we used a solder-able protoboard to manufacture the circuit. Electronic components were connected via a combination of wires and solder bridges between conductive holes. Pin assignments for wiring the Arduino microcontroller board, motor driver, TEC and other components are illustrated in Fig S1. This circuit includes peripherals such as an LCD screen, button controls, and LED indicator lights to fine-tune device functionality. See the operation manual (File S2) for a description of these peripherals and how to use them. The circuit diagram (Fig S1) was drawn with Fritzing (fritzing.org).

#### Arduino sketch for real-time temperature acquisition and control

We employed the programmable Arduino microcontroller to calculate appropriate pulse-width modulated heating output according to set point input. We developed the Arduino sketch (program) in part from existing libraries released by the Arduino community. The sketch was written with Arduino IDE software (version 1.8.1, arduino.cc/en/Main/Software) and loaded via the microcontroller USB port. The sketch is available as supplemental file Script S1.

The sketch was written with the goal of being untethered to a computer, since data collection is not the device’s primary purpose. Temperature data is read in every half second from the thermistor and fed through a proportional integral derivative (PID) loop to determine the power required by the TEC to achieve and maintain the user-defined set point temperature. As the device is powered on, an initial arbitrary set point is displayed on the LCD screen, but the device will not heat or cool until a user manually sets a new set point with the pushbuttons. Once the user specifies the desired set point by releasing the pushbutton, the device begins running the PID function and applying current to the TEC. An LED indicates the intensity of the effort required to heat or cool the heatsink. To retrieve temperature data recorded by the device, we wrote a python script, available as supplemental file Script S2.

#### Temperature detection

To read temperature, the devices employs a 10,000Ω thermistor, which is a type of resistor whose resistance is dependent on temperature. In addition to using readings for real-time feedback, the device records readings every half second and stores them in an array, eventually purging the oldest record for every new one added. To examine device performance, we collected data after programming the device to achieve various setpoint temperatures from various starting temperatures. The conversion of temperature-dependent resistance through the thermistor into temperature is given by the Steinhart-Hart equation:

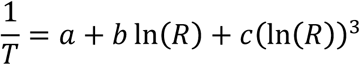

where *T* is temperature in degrees Kelvin, *R* is resistance through the thermistor, and *a*, *b*, and *c* are thermistor-specific coefficients. We then plotted the thermistor temperature readings in degrees Celsius over time to identify the speed at which the device achieved temperature setpoints.

### Characterization of *par-2* temperature sensitive mutations

The temperature sensitive *C. elegans* mutant strain EU1327 *par-2*(*or640* ts) [10] was acquired from the Caenorhabditis Genetics Center (CGC) at the University of Minnesota. The laboratory strain N2 was used as the wild type control. Worms were maintained at the permissive temperature, 15°C, until the experiment was initiated. Temperature shifts to the restrictive temperature, 26°C, were induced as either “long” or “short” upshifts [10]. For the long upshift, late stage L4s were transferred to a 26°C incubator and dissected as young adults to retrieve embryos 4hrs later. For the short upshift, young gravid hermaphrodites were picked to a pre-warmed plate in under 30sec, a timer was started, and the plate was incubated for 5min at 26°C before embryos were retrieved by dissection. In both cases, we used the temperature control device to maintain the restrictive temperature through dissection, slide mounting, and transfer to the heated microscope stage for image capture. Embryos were dissected from gravid adults and agar mounted by standard protocol [11]. Developing embryos were recorded via DIC microscopy using a 40X 0.75AP Nikon air objective and Volocity (PerkinElmer) software.

Once slide-mounted, embryos were screened on the imaging platform and those at the two-cell stage or earlier were recorded by video. All long upshift embryos experienced the restrictive temperature over their entire development, but we timed the short upshift experiments so that our dataset would include embryos that were shifted to the restrictive temperature before, during and just after pronuclear migration. To verify that the *par-2*(*or640* ts) allele is indeed short-acting and to resolve the points at which PAR-2 is required for normal development, we determined when each embryo experienced the shift to the restrictive temperature by projecting backwards from the observed developmental stages in the video using the elapsed time on the timer. We derived quantitative measurements for two phenotypes, the size ratio of the P_1_ and AB cells, given by their long axis lengths, in the two-cell embryo, and the time interval between the division of AB and P_1_ in the transition from the two- to four-cell embryo. Cell sizes were determined from calibrated, still images in Image Pro 10 (Media Cybernetics).

### Identification of thermal ceiling for *Caenorhabditis* embryos

The following *Caenorhabditis* strains, acquired from the CGC, were used in the study: AF16 (*C. briggsae*), EM464 (*C. remanei*), JU1199 (*C. afra*), JU1373 (*C. tropicalis*), JU1667 (*C. monodelphis*), N2 (*C. elegans*). Animals were reared at 20°C under standard nematode culture conditions [5] and shifted to the experimental temperature at the L4 stage of the parental generation using an incubator. Embryos were retrieved, slide-mounted, and imaged as above, using the temperature control device to maintain the experimental temperature during dissection. We selected experimental temperatures to assay lethality in each strain based on our prior observations. The recorded videos were examined frame by frame to identify time of death. To analyze survival within each temperature treatment, we used the *survfit* function in the R package *survival* [12, 13] and visualized results using the package *survminer* [14]. To predict the LT50 temperature at which 50% of embryos should die, we regressed the number of observed dead and surviving embryos onto temperature using the *glm* function with a binomial error structure in R, then used the *dose.p* function in the package *MASS* [15].

## RESULTS

### Device performance

We tested the ability of the device to achieve and stably maintain temperatures between 15°C and 31°C, which represent typical low and high ends of experimental temperature ranges for systems like *Drosophila* and *Caenorhabditis*. For each of four tested temperatures, which include warming to above and cooling to below room temperature, the device stabilized near the set point within a few minutes and continued to converge to the set point over time, eventually achieving stability +/− 0.1°C for the duration of the time data were collected (Fig 2).

**Fig 2.**
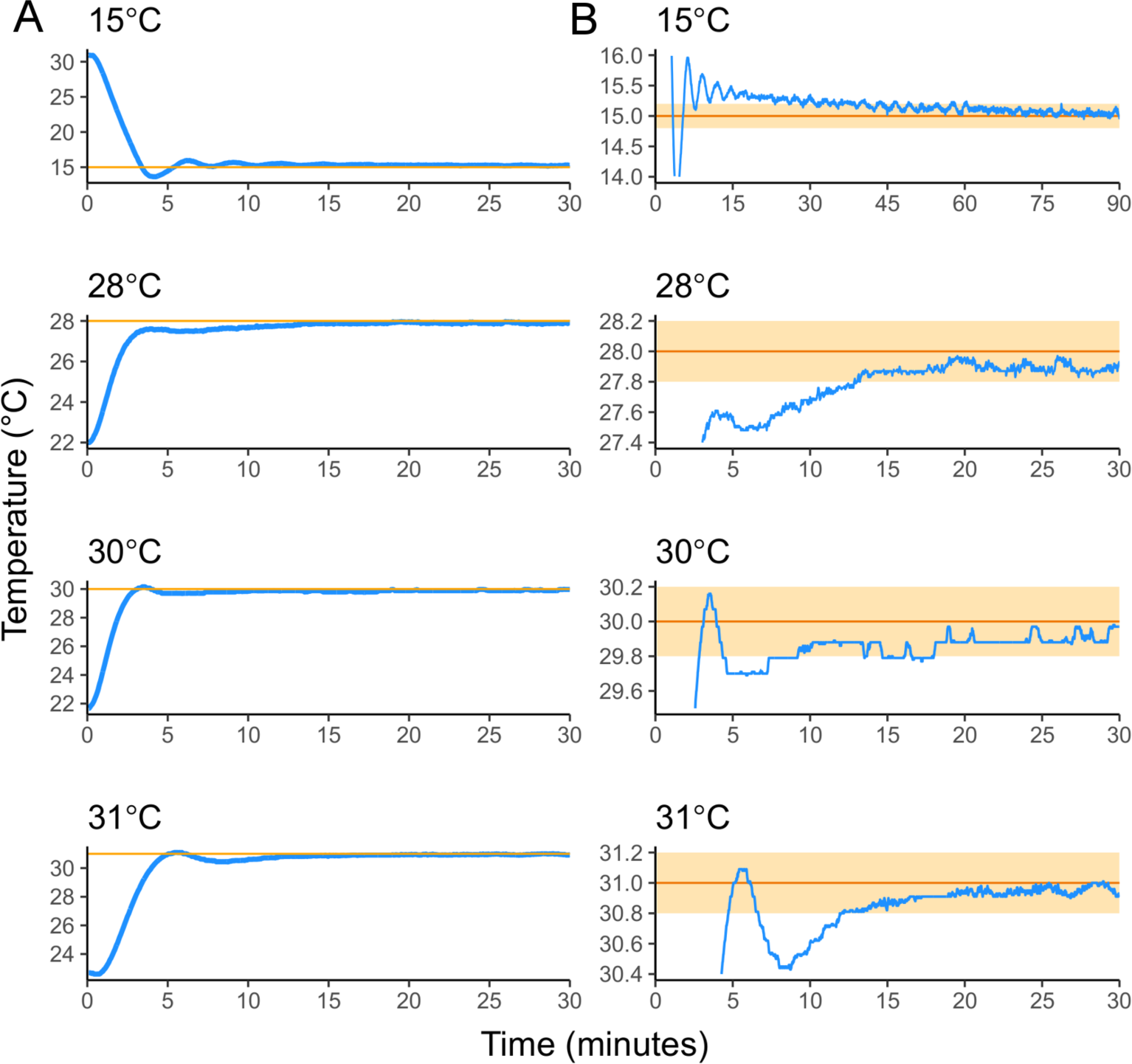
Temperature curves showing how quickly the device achieves set point temperatures. (A) The device achieved stability within 1°C of all set point temperatures (orange lines) within five minutes, including the transition to 15°C from 32°C. (B) Device temperature will continue to converge to the set point over time, achieving stability +/− 0.2°C within 15 minutes and +/− 0.1°C within 30 minutes for the warmer set points. Cooling to 15°C took longer, and was achieved to +/− 0.2°C in about an hour.

### Re-assessment of a fast-acting *par-2* temperature-sensitive allele

One application for this device is to enable or improve manipulation of temperature sensitive mutations. A major challenge to studying essential genes in model systems is that their function is required for viability. Temperature sensitive alleles offer conditional, experimentally induced interruption of function: they typically contain an amino acid variant that destabilizes the protein at a warm, “restrictive” temperature but allows wild type function at a cool, “permissive” temperature. Temperature sensitive alleles were foundational tools in the elucidation of the genetic pathways that govern the first cell divisions of the *C. elegans* embryo, a primary model for understanding cell polarity in animals, but the difficulty of generating and identifying conditional mutations has limited the pool of available alleles in this system [8, 16, 17]. However, recent efforts, including improved methods for mapping mutations, have contributed many new temperature sensitive alleles and a resurged interest in their utility [18–20]. Of particular interest is the discovery that many temperature sensitive alleles are fast acting, inducing the mutant phenotype only a minute or minutes after the switch from permissive to restrictive temperature [10, 18]. Fast action offers experimental power for testing when genes function, but also demands a high level of experimental control, to avoid exposure to restrictive temperatures during sample preparation or even during routine animal husbandry.

Previous work evaluating 24 *C. elegans* temperature sensitive, embryonic lethal mutations found that over half were fast acting, a potential underestimate that the authors suggest may be overcome with better temperature control during embryo mounting [10]. Each of three tested *par-2* mutations were relatively fast-acting, showing the mutant phenotypes of symmetry in the two cell embryo and synchrony in the two-to-four cell division. However, the observed penetrance was incomplete; for example, the authors observed only 3 out of 5 embryos carrying the *par-2*(*or640* ts) allele exhibited mutant phenotypes when the restrictive temperature was applied shortly before phenotyping, compared to 6 out of 6 when the upshift occurred hours in advance. Moreover, because PAR-2 appears to be required prior to pronuclear migration, the short upshift period itself was 30min, limiting precise identification of the critical activity window.

Here, we applied consistent restrictive temperature to *par-2*(*or640* ts) embryos, using the temperature control device to maintain the thermal micro-environment from the initiation of the temperature upshift through embryo dissection, slide mounting and imaging. Some embryos were exposed to the restrictive temperature in a long upshift of 4hrs before screening; others were exposed to a short upshift of several minutes, occurring before, during, or just after pronuclear migration. Wild type embryos show asymmetry in cell size at the two-cell stage, and asynchronous timing in the division of the AB and P_1_ cells in the transition from two- to four-cell stage. We screened the *par-2*(*or640* ts) embryos for loss of asymmetry and loss of asynchrony and observed complete penetrance of both mutant phenotypes in the long upshift embryos (Fig 3). For the short upshift embryos, penetrance of the first phenotype depended upon when the embryo was exposed to the restrictive temperature: embryos upshifted 9min or more before pronuclear migration exhibited near complete symmetry (average two-cell ratio = 0.93, standard deviation = 0.04), whereas those upshifted later in their development showed an intermediate phenotype (ave=0.76, sd=0.06), with some falling into the range observed for wild type (ave=0.64, sd=0.02). Short upshift embryos showed near complete penetrance in the second phenotype, as all embryos exhibited loss of asynchrony, but expressivity was stronger in embryos that were upshifted earlier: those exposed to the restrictive temperature 10min or more before pronuclear migration exhibited perfectly synchronous division from the two- to four-cell stage (average time between cell divisions = 0.0 seconds, standard deviation = 0.0), but most embryos fall into a range of intermediate phenotype (ave=56.33, sd=21.53), clearly asynchronous but not quite wild type (ave=91.2, sd=1.30) (Fig 3). Notably, *par-2*(*or640* ts) embryos with approximately wild type asymmetry at the two-cell stage went on to show intermediate or extreme synchronicity at the next cell division, demonstrating rapid response to PAR-2 perturbation. We also observed weakened expressivity in the two-cell ratio for two embryos assayed earliest in our attempts at this method, which may have experienced brief (<30sec) exposure to room temperature during handling. We hypothesize that potentially even a small departure away from the restrictive temperature may permit rapid re-stabilization of some molecules, affecting function.

**Fig 3.**
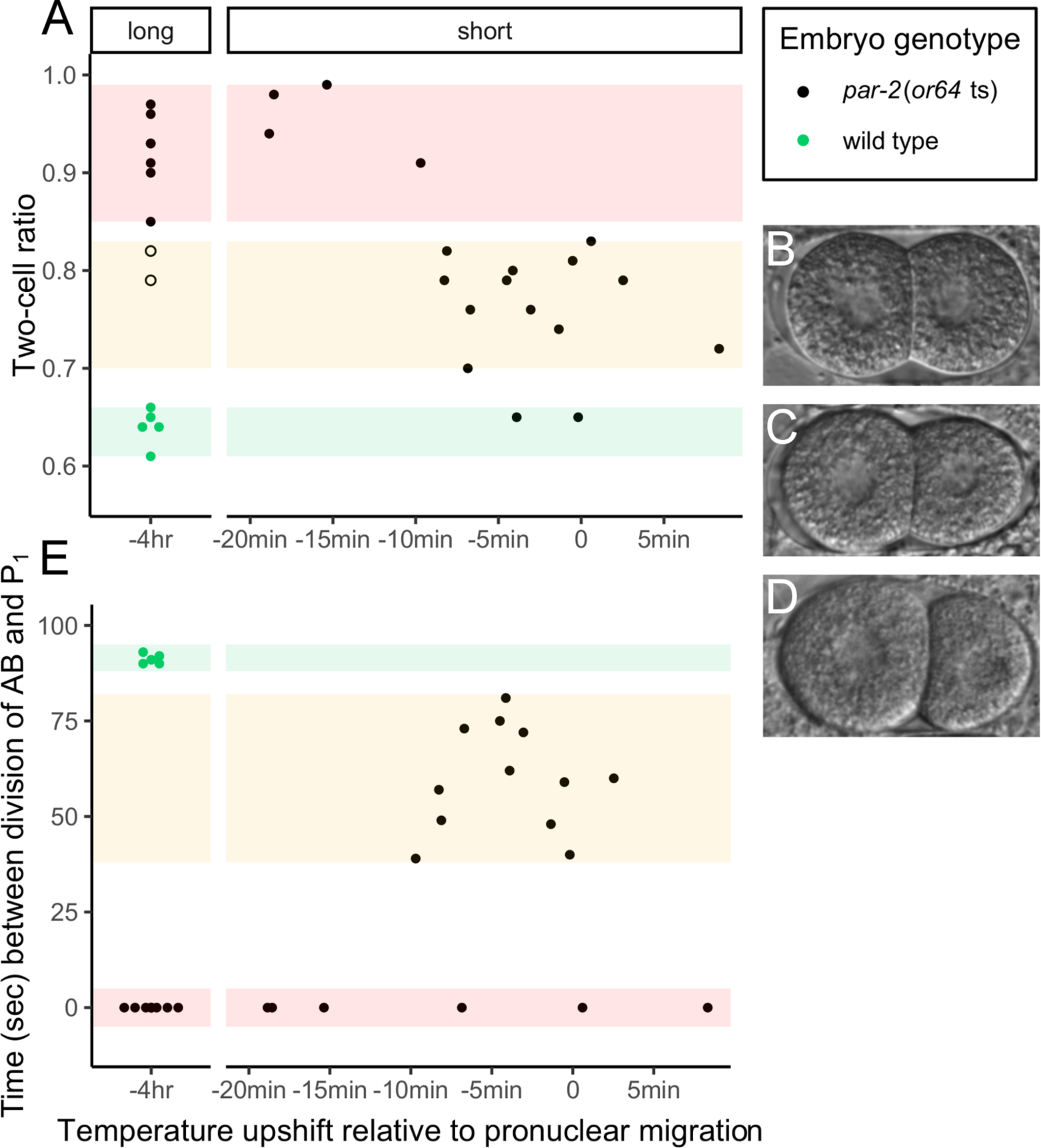
Penetrance and expressivity of a *par-2* temperature sensitive mutation. (A) At the two-cell stage, *par-2*(*or640* ts) embryos exposed to the restrictive temperature in a long upshift (4hrs) or in a short upshift begun at least 9min before pronuclear migration exhibit substantial loss of asymmetry. The range of ratio values for wild type embryos is highlighted in green, and the so-described “extreme” and “intermediate” mutant phenotypes are in red and yellow. Open circles represent two embryos that may have experienced brief exposure to non-restrictive temperature. (B) Two-cell *par-2*(*or640* ts) embryo with extreme loss of asymmetry (ratio=0.91). (C) Two-cell *par-2*(*or640* ts) embryo with intermediate loss of asymmetry (ratio=0.81). (D) Wild type embryo (ratio=0.64). (E) In the transition from two to four cells, embryos upshifted 10min or more before pronuclear migration show complete synchrony in the timing of the division of AB and P_1_ cells, whereas those upshifted later more often show an intermediate time interval between cell divisions.

In both of the *par-2* phenotypes we examined, we observed a roughly stepwise pattern, in which embryos show either extreme or intermediate loss of asymmetry/asynchrony, determined by a timepoint about 9-10min before pronuclear migration. Additional observations may be needed to confirm whether this phenomenon is discrete, or if the traits are actually continuous. Overall, we draw two conclusions from these results: that improved control of the thermal micro-environment can substantially improve phenotypic penetrance, and that the *par-2*(*or640* ts) allele is indeed fast-acting.

### *Caenorhabditis* species show variable thermal ceilings

While poikilotherms can exhibit dramatic developmental plasticity in response to their thermal environment, and may tolerate a wide range in temperature, the molecular and cellular mechanisms that mediate temperature-dependent development may nevertheless be finely tuned and adapted [21, 22]. This is evidenced by the many observations that organisms exhibit variation in critical temperatures for fitness, tolerance and preference, both within and between species. For example, relative to *C. elegans*, *C. briggsae* fitness is optimized at a higher temperature [23, 24] and *C. briggsae* embryos first exhibit signs of heat stress at a higher thermal ceiling [6], though both species exhibit heritable variation for thermal preference or thermal-mediated fecundity [25–27]. Likewise, across the genus *Drosophila*, species show uniform scaling in the timing of landmark developmental events, but with distinct rates with regard to temperature [3]. Consequently, studies of comparative development are challenged not only to carefully control the thermal environment of experimental specimens, but to select appropriate temperatures, which will vary by strain or species [3].

In other research, we sought to compare early embryonic development across the genus *Caenorhabditis*, and uncover potential divergence in the molecular mechanisms that govern it, by exaggerating morphological differences using heat stress. To that end, first we needed to identify species-specific thermal ceilings for embryonic viability. For six species, we recorded embryonic development from the one- through four-cell stage under thermal stress and determined the time of death (Fig 4). For *C. monodelphis, C. remanei* and *C. tropicalis*, we assessed viability at multiple temperatures, then fit the data to a linear model to predict the “LT50”, or lethal temperature at which 50% of embryos fail to survive; for *C. elegans*, *C. briggsae* and *C. afra*, we followed up on prior observations by assessing viability at a single temperature.

**Fig 4.**
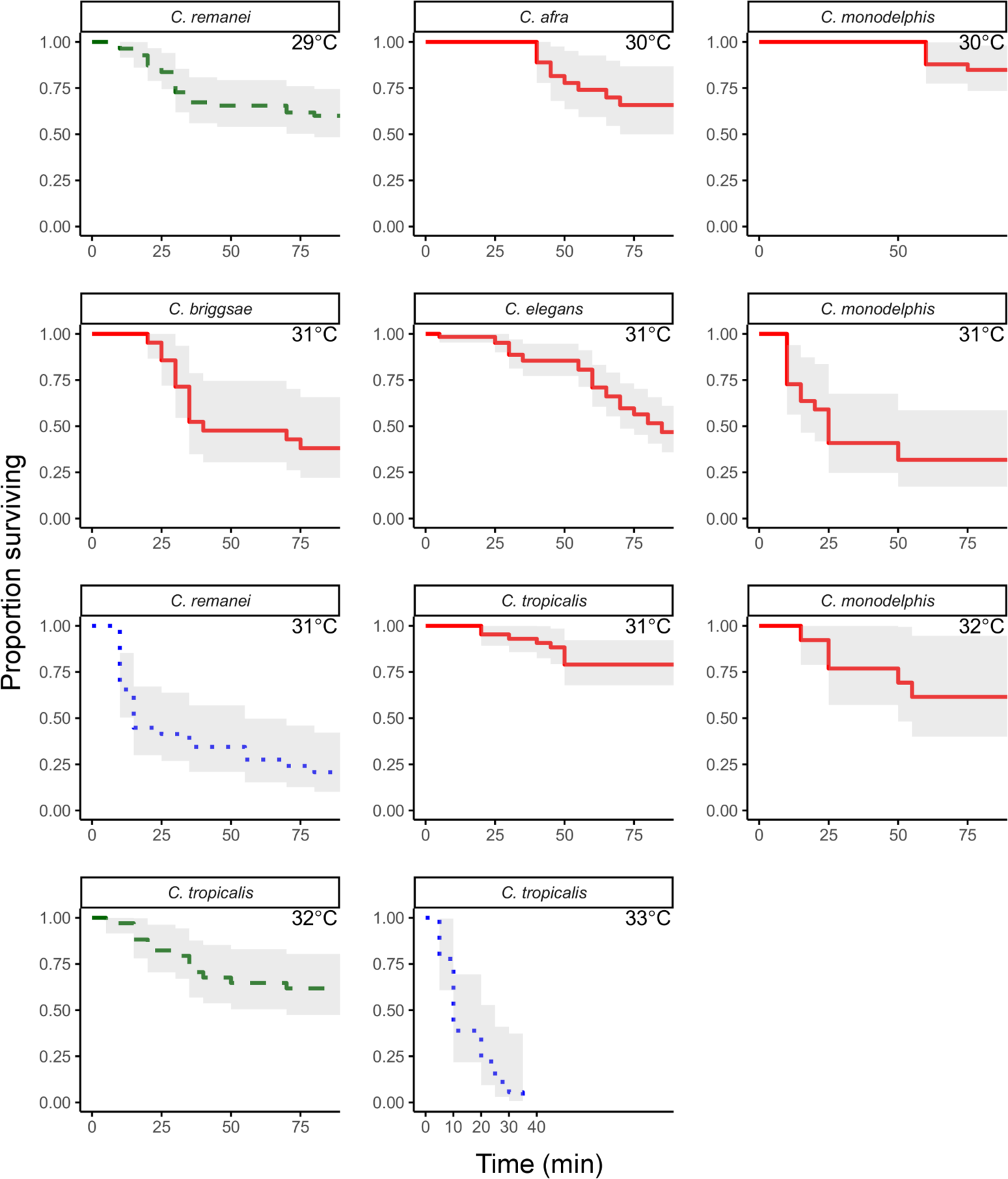
Survivorship curves for *Caenorhabditis* species. Despite extreme conservation in morphological development, *Caenorhabditis* species demonstrate distinct differences in ability to survive heat stress. For example, nearly 50% of embryos from the temperate species *C. remanei* die within 90min when reared at 29°C, but embryos of the tropical species *C. tropicalis* must be exposed to 32°C to exhibit the same survivorship pattern (green dashed lines). However, at just one degree higher, 33°C, *C. tropicalis* embryos show severe and rapid lethality, while a two degree increase for *C. remanei*, to 31°C, does not induce as dramatic an increase in lethality (blue dotted lines), suggesting a potential biophysical ceiling for adaptation to high heat.

We found that tropically distributed *C. tropicalis* demonstrated the highest lethal temperature, with a predicted 50% embryonic lethality at 31.95°C (standard error ± 0.14), and the temperate species *C. remanei* the lowest (T50 = 29.41°C ± 0.14). Perhaps unexpectedly, we identified a relatively high lethal temperature for *C. monodelphis* (T50 = 31.36°C ± 0.69), a species distantly related to *C. elegans* (and all other species in the genus) that has been sampled from northern Europe [28]. Although *C. briggsae* fitness is known to be optimized at a higher temperature than *C. elegans* [23, 24], our *C. briggsae* embryos reared at 31°C did not survive at a higher proportion than *C. elegans* embryos (33% and 44%, respectively). This may indicate that fitness-optimized temperatures will not necessarily co-vary with ceilings for thermal tolerance under high stress conditions. Consistent with this and with other observations that *C. elegans* embryos exhibit signs of heat stress at lower temperatures than *C. briggsae* [6], we did observe twice as many cell division aberrations in *C. elegans* embryos reared at 31°C relative to *C. briggsae* (data not shown). We also note that thermal ceilings for one- through four-cell embryos may be higher than those of later stage embryos, since *C. elegans* embryos show developmental arrest at later stages at lower temperatures [29].

## DISCUSSION

We describe an easy to use, portable temperature control device that can achieve a setpoint temperature above or below ambient temperature in minutes, and stably maintain that temperature to within 0.1°C. We propose that this device may be useful to biologists that require precise control of specimen temperature outside of an incubator, for example when live samples must be manipulated in preparation for microscopy imaging. Of course, this device or versions of it may be of use outside of biology research, for any application that requires simple manipulation of micro-environmental temperature.

In the biological context, our device might be compared to commercially available microfluidics systems specifically designed to control micro-environmental temperature in the context of *C. elegans* microscopy. However, unlike thin, fluidics-based devices constructed of transparent polymers, which can shift temperature in seconds and simultaneously support image capture on compound or confocal microscopes [30], our device is not suited for inducing rapid temperature change or for use under high magnification microscopy. Rather, it provides complementary functionality, such as thermal control during the preparation of samples, which may then be loaded into such devices, conventionally slide-mounted and imaged on a temperature-controlled stage, or otherwise processed. At a materials cost of several hundred USD, it is also less than 1/30 or 1/50 the price of the two fluidics-based systems currently developed [30].

## Supporting information

Dataset S1 par-2 phenotypes

Figure S1 circuit diagram

File S1 device partslist

Script S1 arduino sketch

Dataset S2 Cae survivorship

File S2 operation manual

Script S2 data transfer

## ACKNOWLEDGMENTS

This research was funded by NIH grant R35 GM119744 to ABP. Worm strains were provided by the Caenorhabditis Genetics Center, which is funded by NIH Office of Research Infrastructure Programs (P40 OD010440). We thank the Parker H. Petit Institute for Bioengineering and Bioscience at the Georgia Institute of Technology for the use of their shared equipment, services and expertise, in particular the valued assistance of graphic designer Tim Whelan for his contributions to the technical drawing and Dr. Aaron Lifland in the Optical Microscopy Core for his help with imaging.

## SUPPORTING INFORMATION

**S1 Figure. Circuit diagram.** Orange lines indicate positive wires, black indicate ground, and teal indicate Arduino input/output.

**S1 File. Partslist for device.**

**S2 File. Operation manual for using the device.**

**S1 Script. Script for Arduino sketch.**

**S2 Script. Python script for data transfer from device to computer.**

**S1 Dataset. *C. elegans par-2* ts data.** The “ratio” column provides cell length ratio in the two-cell embryo. The “delta” column provides the time, in seconds, between the division of the AB and P_1_ cells in the transition to the four-cell embryo. The “time_PM” provides the time, in seconds, at which pronuclear migration occurred relative to the temperature upshift.

**S2 Dataset. *Caenorhabditis* embryonic lethality data.** The “status” column indicates whether the embryo survived (0) or died (1). For surviving embryos, the “time_of_death” column will give the time the video ended. Time is given in minutes.

