## Supplementary figures and images for "A portable, low-cost device for precise control of specimen temperature under stereomicroscopes"

### Figure S1 circuit diagram

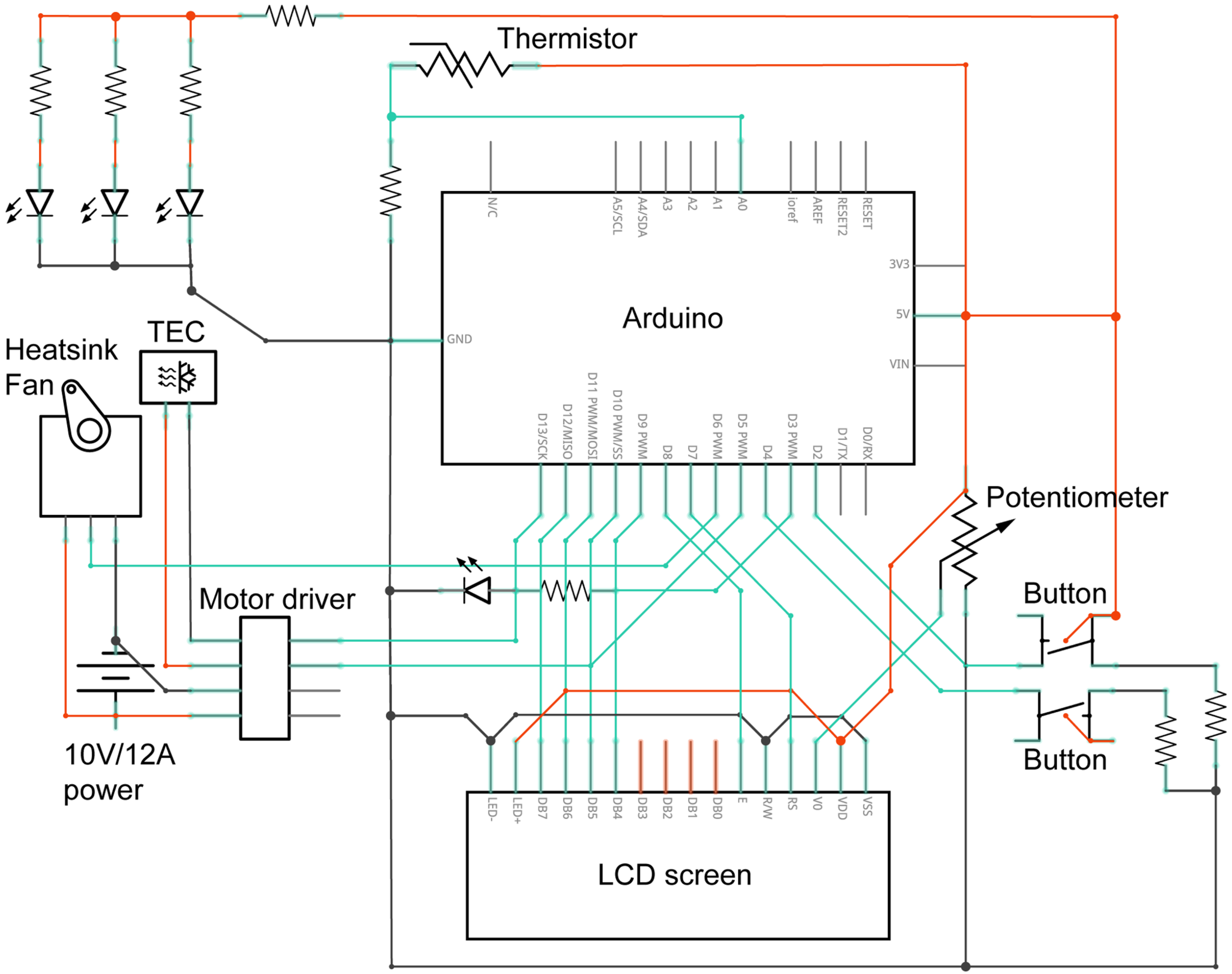
