## Supplementary material for "A portable, low-cost device for precise control of specimen temperature under stereomicroscopes": File S1 device partslist

| Part | Cat. / Part No. | Quant. | Source | Cost |
|---|---|---|---|---|
| TEC (Peltier) | 64975-502 | 1 | Laird Technologies | $67.63 |
| thermistor | SA2F-TH-44031-40 | 1 | Omega | $60.00 |
| Arduino starter kit | Arduino Starter Kit - ENG | 1 | Arduino.com | $99.90 |
| Arduino Uno Rev. 3 | | 1 | | |
| Alphanumeric LCD, 16x2 characters | | 1 | | |
| Breadboard, 400 points | | 1 | | |
| Solid core jumper wires, 22 gauge | | 70 | | |
| Wooden base | | 1 | | |
| 220 Ohm resistor | | 3 | | |
| Pushbuttons | | 2 | | |
| Potentiometer, 10k Ohms | | 1 | | |
| LED, bright white | | 1 | | |
| motor driver | SHIELD-MD10 | 1 | Cytron | $13.78 |
| protoshield | PROTO-01 | 1 | OSEPP | $6.99 |
| bottom heatsink | Heatsink R18 | 1 | Dynatron | $29.59 |
| top heatsink | - | 1 | custom made | - |
| thermal paste | XMS99 | 2 | Arctic Silver 5 | 11.99 |
| Arduino power supply | 101-60-182 | 1 | SainSmart | $7.99 |
| mirror (square 3x3”) | ASIN B004HGODQU | 1 | Amazon.com | $5.58 |
| etching cream | 15-0200 | 1 | Armour Products | $21.09 |
| TEC power supply (12V/10A output) | ASIN B00Z9X4GLW | 1 | Amazon.com | $17.99 |
| power supply cable (5.5mm x 2.1mm) | ASIN B072BXB2Y8 | 1 | Amazon.com | $6.98 |
| | | | | $349.51 |
