## Supplementary material for "A portable, low-cost device for precise control of specimen temperature under stereomicroscopes": Script S1 arduino sketch

/********************************************************
 * TEC control
 ********************************************************/

#include <LiquidCrystal.h>
#include <math.h>
#include <PID_v1.h>

unsigned long timeMilli;
//Define Variables we'll be connecting to
double Setpoint = 22;     //set the original Setpoint = 20C
double Input, Output;
//Define the aggressive and conservative Tuning Parameters
double aggKp = 9,   aggKi = 0.05,  aggKd = 1;
double consKp = 100, consKi = 0.25, consKd = 4;

//define button-controlled setpoint adjustments
float tempAdjNeg, tempAdjPos, LEDtemp;
int minusPushCounter = 0; //counter for number of button presses
int minusState = 0;       //current state of button
int lastMinusState = 0;   //previous state of button
int plusPushCounter = 0; //counter for number of button presses
int plusState = 0;       //current state of button
int lastPlusState = 0;   //previous state of button

int fanOutput = 0;

PID myPID(&Input, &Output, &Setpoint, aggKp, aggKi, aggKd, DIRECT);

#define ThermistorPIN A0
#define minusPIN 2
#define LEDpin 3
#define plusPIN 4
#define PeltierPIN 5                      // PWM Pin 5
#define dirPIN 13                         // determines direction of the peltier
#define fanPIN 6

LiquidCrystal lcd(7, 8, 9, 10, 11, 12);   // Pins for LCD display

float vcc = 4.91;                       // only used for display purposes, if used
// set to the measured Vcc. <- voltage of power supply
float pad = 9850;                       // balance/pad resistor value, set this to
// the measured resistance of your pad resistor
float thermr = 10000;                   // thermistor nominal resistance

int Vo;
float R1 = 10000;
float logR2, R2, T, Tc, Tf;
float c1 = 1.009249522e-03, c2 = 2.378405444e-04, c3 = 2.019202697e-07;

const int numReadings = 10;
double readings[numReadings];      // the readings from the analog input
int readIndex = 0;              // the index of the current reading
double total = 0;                  // the running total

int pressed = 0;              // determines whether a button has ever been pressed

float Thermistor(int RawADC)
{
  long Resistance;
  float Temp;  // Dual-Purpose variable to save space.

  Resistance = pad * ((1024.0 / RawADC) - 1);
  Temp = log(Resistance);    // Saving the Log(resistance) so not to calculate  it 4 times later
  Temp = 1 / (0.001129148 + (0.000234125 * Temp) + (0.0000000876741 * Temp * Temp * Temp));
  Temp = Temp - 273.15;  // Convert Kelvin to Celsius
  //Temp = (Temp * 9.0)/5.0+32.0; //C to F
  return Temp;                                      // Return the Temperature
}

void setup()
{
  pinMode(plusPIN, INPUT);
  pinMode(minusPIN, INPUT);
  pinMode(LEDpin, OUTPUT);
  pinMode(PeltierPIN, OUTPUT);
  pinMode(dirPIN, OUTPUT);
  //code from thermistor reader:
  Serial.begin(115200);

  //Code from PID: initialize the variables we're linked to
  Input = Thermistor(analogRead(ThermistorPIN));

  // initialize all the readings to 0:
  for (int thisReading = 0; thisReading < numReadings; thisReading++)
  {
    readings[thisReading] = 0;
  }

  //turn the PID on
  myPID.SetOutputLimits(-255, 255); //tells PID to range output from -255 to 255
  myPID.SetMode(MANUAL);            //turns off PID until a button is pressed (below)

  //turns on the lcd display
  lcd.begin(16, 2);

}

void loop()
{
  //read in input buttons
  plusState = digitalRead(plusPIN);
  minusState = digitalRead(minusPIN);

  //read in resistance across thermistor; convert to temp
  float temp;
  //  temp = Thermistor(analogRead(ThermistorPIN));       // read ADC and  convert it to Celsius
  // subtract the last reading:
  total = total - readings[readIndex];
  // read from the sensor:
  readings[readIndex] = Thermistor(analogRead(ThermistorPIN));
  // add the reading to the total:
  total = total + readings[readIndex];
  // advance to the next position in the array:
  readIndex = readIndex + 1;

  // if we're at the end of the array...
  if (readIndex >= numReadings)
  {
    // ...wrap around to the beginning:
    readIndex = 0;
  }

  // calculate the average:
  temp = total / numReadings;
  // send it to the computer as ASCII digits
  delay(1);        // delay in between reads for stability


  Input = temp;

  // compare buttonStates to their previous state
  if (minusState != lastMinusState)
  {
    // if the state has changed, increment the counter
    if (minusState == HIGH)
    {
      // if the current state is HIGH then the button
      // went from off to on:
      //minusPushCounter++;
      tempAdjNeg = 0.5;
      pressed = 1;
    }
    else
    {
      tempAdjNeg = 0;
    }
    // Delay a little bit to avoid bouncing
    delay(100);
  }
  if (plusState != lastPlusState)
  {
    // if the state has changed, increment the counter
    if (plusState == HIGH)
    {
      // if the current state is HIGH then the button
      // went from off to on:
      //plusPushCounter++;
      tempAdjPos = 0.5;
      pressed = 1;
    }
    else
    {
      tempAdjPos = 0;
    }
    // Delay a little bit to avoid bouncing
    delay(100);
  }
     
  if (tempAdjPos == 0 & tempAdjNeg == 0 & pressed == 1)    //turns PID on AUTO after button is released
  {
    myPID.SetMode(AUTOMATIC);
  }

  //this configuration should make it so that Setpoint adjusts only based on button input
  Setpoint = Setpoint + tempAdjPos - tempAdjNeg;
  //Input = temp;

  //adaptive tuning parameters for PID
//  double gap = abs(Setpoint - Input); //distance away from setpoint
//  myPID.SetTunings(aggKp, aggKi, aggKd);
//  if (gap < 2)
//  {
//    myPID.SetTunings(consKp, consKi, consKd);
//  }
  if (Setpoint < temp)
  {
    myPID.SetTunings(consKp, consKi, consKd);
  }
  if (Setpoint >= temp)
  {
    myPID.SetTunings(aggKp, aggKi, aggKd);
  }
  myPID.Compute();

  if (Output < 0)
  {
    digitalWrite(dirPIN, HIGH);       //switch peltier to cooling mode
    Output = Output * -1;
  }
  else
  {
    digitalWrite(dirPIN, LOW);       //keeps it on heating mode
  }

  fanOutput = Output * 0.75;
  analogWrite(PeltierPIN, Output);
  analogWrite(fanPIN, fanOutput);
  analogWrite(LEDpin, Output);

  // set the cursor to (0,0):
  lcd.setCursor(0, 0);
  lcd.print("Set: ");
  lcd.print(Setpoint, 1);
  lcd.print("C");
//  lcd.print(Output);
  //  lcd.print(dirChange);
  // set the cursor to (16,1):
  lcd.setCursor(0, 1);
  lcd.print("Temp: ");
  lcd.print(Input, 1);
  lcd.print("C");
  tempAdjPos = 0;
  tempAdjNeg = 0;
  // clear screen for the next loop:
  delay(500);
  lcd.clear();

  // print information to computer
  timeMilli = millis();     // time since program started
  Serial.print(timeMilli);  //prints time since program started
  Serial.print(", ");
  Serial.print(Setpoint, 2); // print setpoint as dictated by button press
  Serial.print(", ");
  Serial.print(Input, 2); // print temperature after conversion
  Serial.print(", ");
  Serial.print(Output, 2); // print how hard TEC is working to get to setpoint 
  Serial.print(", ");
  Serial.println(fanOutput, 2); // print how hard fan is working to cool off heatsink

}
