## Supplementary material for "A portable, low-cost device for precise control of specimen temperature under stereomicroscopes": File S2 operation manual

### Temperature control device operation manual

#### General operating instructions

- To turn on, plug the two power cords into electrical outlets. Or, use a power strip and use the power strip switch to power up/down the device.
- Once on, the LCD screen will illuminate and should say something like:

```
Set: 22.0C
Temp: 21.7C
```

- The device will not activate until the “Set” temperature has been adjusted.
- To adjust the “Set” temperature, locate the temperature set point + and − buttons on the top of the device, located towards the middle of the circuit board. If unlabeled, the + button is above the – button (with LCD screen facing you) and they are very close together.
- Press and hold either button to adjust the settings. The input values will increase/decrease in 0.5˚C increments.
- Now the device will bring the stage up to temperature and maintain within 0.1˚C. In 5min, the temperature should be stable within 1˚C of the set point. By 30min, the device stage should be completely stable around the set point temperature, assuming typical environmental conditions.
- The LED indicator will display how much effort is being put into achieving/maintaining the stage temperature. It should be bright initially, and dim significantly as temperature stability is achieved.
- If needed, a reset button is provided just below the LED indicator. Press this button to reset the temperature settings. Note this action clears the PID parameters, so it is unwise to use this unless you are switching between heating and cooling.

#### Troubleshooting

*Is the LCD screen spitting out gibberish and a flurry of non-standard characters?*

There is likely a connection issue between one (or more) wires connecting the arduino to the LCD screen. Try jiggling the screen/wires and reset.

*Arduino turns on and functions, but the temperature doesn’t respond and you don’t hear the fan.*

Check the motor driver shield. Is it blinking or failing to turn on, as indicated by the running LED and/or the A & B LEDs? This is likely due to a bad power supply.

*Is the LCD/arduino flickering and/or not powering on properly?*

This is also likely due to a bad power supply.

#### Interfacing with the device

**Ports**

- Arduino USB port

  *Located on arduino. Use to interface with a computer, e.g. updating code or collecting data. Not required for normal device function.*

- Power port

  *Located on arduino (bottom board)*

- Power port

  *Located between motor control shield (middle board) and protoshield (top board)*

**Inputs**

- Thermistor

  *Collects temperature data from the working surface; no human interface necessary.*

- Reset button

  *Located on top right of board (with LCD screen facing you). Use to reset PID parameters and the user-defined temperature set point.*

- Temperature + button

  *Located near middle of board, just above – button.*

- Temperature – button

  *Located near middle of board, just below + button.*

- Potentiometer

  *Located to the left of the temperature control buttons. Adjusts contrast on the LCD screen.*

**Outputs**

- LCD screen

  *Displays current user-defined temperature set point (top line) and current temperature as measured by thermistor (bottom line).*

- LED indicator, PID output

  *Located on the top right of the top board. The brightness of this LED displays the current PID output being sent to the stage to heat or cool. Full brightness indicates a 100% effort, suggesting that the device is working hard to achieve/maintain temperature.*

- LED indicator, motor control output

  *Located on the far side of the motor control board, on the far right. Indicates whether the motor control board is functioning properly. Used for troubleshooting board functioning.*

- LED indicators, motor control direction

  *Located just to the left of motor control indicator, labeled A and B. These indicate the direction of electricity flow through the board. When A is lit, the stage is cooling. When B is lit, stage is heating.*
